# Combination of CG methylation and Chromomethylase 2-mediated RdDM-independent CHH methylation is required for chromosome-specific rRNA gene silencing

**DOI:** 10.1101/2023.02.03.526984

**Authors:** Navinchandra V. Puppala, Ramya Tammineni, Gargi P. Saradadevi, Gireesha Mohannath

**Affiliations:** Institute for Developmental Biology and Neurobiology, Johannes Gutenberg University Mainz, Biozentrum I, Mainz, Germany; Department of Biological Sciences, Birla Institute of Technology and Science-Pilani, Hyderabad campus, Hyderabad, Telangana, India; Stem Cells Division, SKAN Research Trust, St. John’s Research Institute, Bengaluru, Karnataka, India

**Keywords:** Ribosomal RNA (rRNA), rDNA promoter, Nucleolus organizer region (NOR), Gene silencing, DNA cytosine methylation, Chromomethylase 2 (CMT2), DDM1, MET1

## Abstract

*Arabidopsis thaliana* has two nucleolus organizer regions (NORs), consisting of long tandem arrays of 45S rRNA genes on chromosomes 2 (*NOR2*) and 4 (*NOR4*). The rRNA gene subtypes mapping to *NOR2* are mostly silenced during development, whereas most of those mapping to *NOR4* are active. Cytosine methylation of promoters plays a significant role in rRNA gene silencing, but recent findings demonstrate that gene body methylation also play a key role in gene silencing. Previous studies have demonstrated that CG methylation plays a role in rRNA gene silencing. However, it remains to be determined whether Chromomethylase 2 (CMT2)-mediated, RdDM-independent CHH methylation plays any role in rRNA gene silencing. To find out the relative importance of cytosine methylation in each of the sequence contexts (CG, CHG, and CHH, where H=A, T, or C), we studied multiple DNA methylation-deficient *Arabidopsis* mutants. Primarily, we show that the mutants that displayed higher losses of CG or CHH methylation at rDNA loci, display higher degrees of *NOR2* rRNA gene silencing disruption. Using whole genome bisulfite sequencing data, we show that the loss of RdDM-independent CMT2, not the loss of RdDM-dependent DRM2, erases most of the CHH methylation at rDNA loci, while retaining CG methylation levels comparable to those of wild-type Col-0, demonstrating the sufficiency of CHH methylation loss for releasing rRNA gene silencing. Analysis of *Arabidopsis* and tomato NOR sequences revealed that in 45S rRNA gene sequences, cytosines mostly occur in the CHH context, followed by CG and the lowest being in CHG contexts. The predominance of CHH sites is even more pronounced when the rRNA gene promoter and spacer promoter(s) are considered (0-7% CG, 14-25% CHG, and 79-80% CHH), suggesting a mechanism to explain the stronger disruption of *NOR2* gene silencing in mutants that display higher losses of CHH methylation. Our data also reveal a role for gene body methylation in rRNA gene silencing. We show that *NOR2* genes are relatively hypermethylated compared to *NOR4*, consistent with findings from recent studies. By contrast, human rRNA genes have more CG sites than CHH and CHG, correlating with the evolutionary loss of CMTs in mammals. CMTs are plant-specific, which primarily methylate CHG and CHH sites. Our data define a critical role for CMT2-mediated RdDM-independent CHH methylation, in combination with MET1-mediated CG methylation, in rRNA gene silencing, which is entirely mediated by CG methylation in mammals.

## Introduction

In eukaryotes, hundreds to thousands of 45S ribosomal RNA (rRNA) genes are arranged as tandem repeats at chromosomal loci known as nucleolus organizer regions (NORs) (Gerbi, 1986, Long and Dawid, 1980, Viktorovskaya and Schneider, 2015). RNA polymerase I (Pol I) transcribes rRNA genes to produce 45S pre-rRNA transcripts that are then processed into 18S, 5.8S, and 25-28S rRNAs that comprise the catalytic core of ribosomes, which synthesize proteins in cells (Saradadevi et al., 2020, Turowski and Tollervey, 2015, Viktorovskaya and Schneider, 2015). Although eukaryotic genomes contain many rRNA genes, not all of them are transcribed at all times. Instead, the number of genes transcribed varies during development, dictated by the developmental stage-specific need for ribosomes and protein synthesis (Grummt, 2013, Hannan et al., 2013, Moss et al., 2007). rRNA ‘dosage control’ involves selective gene silencing rather than selective gene activation (Chen and Pikaard, 1997, Pikaard et al., 2023). In *Arabidopsis thaliana*, rRNA genes are clustered within two NORs located at shorter arms of chromosomes 2 (*NOR2,* which spans ∼ 5.9 Mb) and 4 (*NOR4,* which spans ∼ 3.9 Mb) (Copenhaver et al., 1995, Copenhaver and Pikaard, 1996b, Copenhaver and Pikaard, 1996a, Fultz et al., 2023). Previous findings in *A. thaliana* demonstrated that the selective silencing of rRNA genes is determined by their chromosomal location, not rRNA gene sequence variation (Chandrasekhara et al., 2016, Mohannath et al., 2016). During development, rRNA genes located on *NOR2* are selectively silenced while most of those on *NOR4* remain active (Chandrasekhara et al., 2016, Mohannath et al., 2016).

Transcriptional gene silencing in eukaryotes often involves cytosine methylation and chromatin condensation (Rhee et al., 2002, Zhang et al., 2006, Zilberman et al., 2007). Recent findings have demonstrated that in addition to gene promoter methylation, gene body methylation also plays a role in gene silencing (Lee et al., 2023a, Shahzad et al., 2025, Williams et al., 2023). Although cytosine methylation has been known to play a role in rRNA gene silencing (Costa-Nunes et al., 2010, Pontvianne et al., 2013, Preuss et al., 2008), the relative role of cytosine methylation in all three sequence contexts (CG, CHG, and CHH, where H=A, C, or T) in selectively silencing *NOR2* rRNA genes remains unclear. In plants, cytosine methylation is deposited by several DNA methyltransferases belonging to different families. *METHYLTRANSFERASE 1* (*MET1*) and *CHROMOMETHYLASE 3* (*CMT3*) mediate virtually all *Arabidopsis* CG and CHG methylation, respectively (Cokus et al., 2008, Lister et al., 2008, Zemach et al., 2013). CHH methylation is directed by small RNAs and mediated by *DOMAINS REARRANGED METHYLTRANSFERASE 2* (*DRM2*) via the RNA-directed DNA methylation (RdDM) pathway, but also by *CHROMOMETHYLASE 2* (*CMT2*) independently of RdDM (Zemach et al., 2013, Law and Jacobsen, 2010, Rymen et al., 2020, Sasaki et al., 2019). *CMT2* and *CMT3* belong to a group of plant-specific chromomethylases (Bewick et al., 2017). In addition to the aforementioned DNA methyltransferases, *DECREASE IN DNA METHYLATION 1* (*DDM1*), a SWI2/SNF2 family chromatin remodeler, acts as a master regulator of DNA methylation (Brzeski and Jerzmanowski, 2003, Lee et al., 2023b, Osakabe et al., 2021). Loss of DDM1 results in genome-wide demethylation at 70% of methylated cytosines in all three sequence contexts (CG, CHG, and CHH) (Jeddeloh et al., 1999, Vongs et al., 1993, Zemach et al., 2013). Recent findings have demonstrated that DDM1-mediated gene body methylation also contributes to silencing of genes involved in defense response (Lee et al., 2023a).

An earlier study investigated the effects of *drm2*, *cmt3,* and *met1* mutants on rRNA gene silencing (Pontvianne et al., 2013). However, these mutants predominantly exhibit a loss of CG or CHG methylation accompanied by a minimal to modest loss of CHH methylation (Cokus et al., 2008, Zemach et al., 2013). Among these, *met1* mutants displayed the maximum disruption of *NOR2* gene silencing, which is attributed mainly to the loss of CG methylation in these mutants. In this study, we analysed *ddm1* and *cmt2* mutants, along with the three previously studied (*drm2*, *cmt3*, and *met1*) DNA methylation-deficient mutants to compare the relative effects of differential loss of CG, CHG, and CHH methylation on the disruption of chromosome-specific *NOR2* rRNA gene silencing. In this mutant panel, we show higher disruption of *NOR2* rRNA gene silencing in the mutants that displayed maximum loss of CHH or CG methylation at rDNA loci. We show that NOR2 genes are relatively hypermethylated than NOR4 genes, consistent with findings from other recent studies (Fultz et al., 2023, Mo et al., 2023).

Using NOR sequences of *Arabidopsis* (Fultz et al., 2023) and tomato (Shirasawa and Ariizumi, 2024), we show that in the 45S rRNA gene sequences, occurrence of cytosines is the highest in the CHH context, followed by CG, and the lowest in the CHG context, providing a putative mechanistic basis for higher *NOR2* derepression observed in *cmt2, ddm1, and met1* mutants that display significantly higher losses of CHH or CG methylation than the other mutants. In contrast to the exclusive involvement of CG methylation in mammalian rRNA gene regulation, our study demonstrates a critical role for CMT2-mediated CHH methylation, in addition to the previously established role for CG methylation, in plant rRNA gene regulation

## Results

### Selective *NOR2* rRNA gene silencing is disrupted in DNA methylation mutants

The rRNA genes in *A. thaliana* are organized into two tandem arrays on NORs, one each on chromosomes 2 (*NOR2*), and 4 (*NOR4*) (Figure 1). In the reference strain Col-0, *NOR2* is 5.5 Mb and *NOR4* is 3.9 Mb, with ∼550 and ∼390 tandem repeats of 45S rRNA genes, respectively (Fultz et al., 2023) (Figure 1). The 18S, 5.8S, and 25S rRNA sequences are identical among 45S rRNA genes, but variation exists in the internal transcribed spacer (ITS) and the external transcribed spacer (ETS) regions. Four distinct variants, named VAR1 through VAR4, had previously been defined based on the length polymorphism in the 3’ ETS region (Pontvianne et al., 2010) (Figure 1B-C). A recent study in the Col-0 ecotype identified 74 rRNA gene subtypes, based on length and sequence variation (Fultz et al., 2023). Nonetheless, the primers that amplify the four rRNA subtypes bind to most of the repeats from both NORs. Of these, VAR1, VAR2, and a subset of VAR3 (VAR3a) lack HindIII sites in the 3’ ETS region, but the remaining variants (VAR3b,c and VAR4) have a HindIII site (Chandrasekhara et al., 2016, Mohannath et al., 2016) (Figure 1C). During development, the rRNA gene variants that map primarily to *NOR2* (VAR1, VAR3a) are selectively silenced as a consequence of dosage control. In contrast, the variants that map primarily to *NOR4* (VAR2, VAR3c, and VAR4) remain active (Figure 1D)(Chandrasekhara et al., 2016).

**Figure 1:**
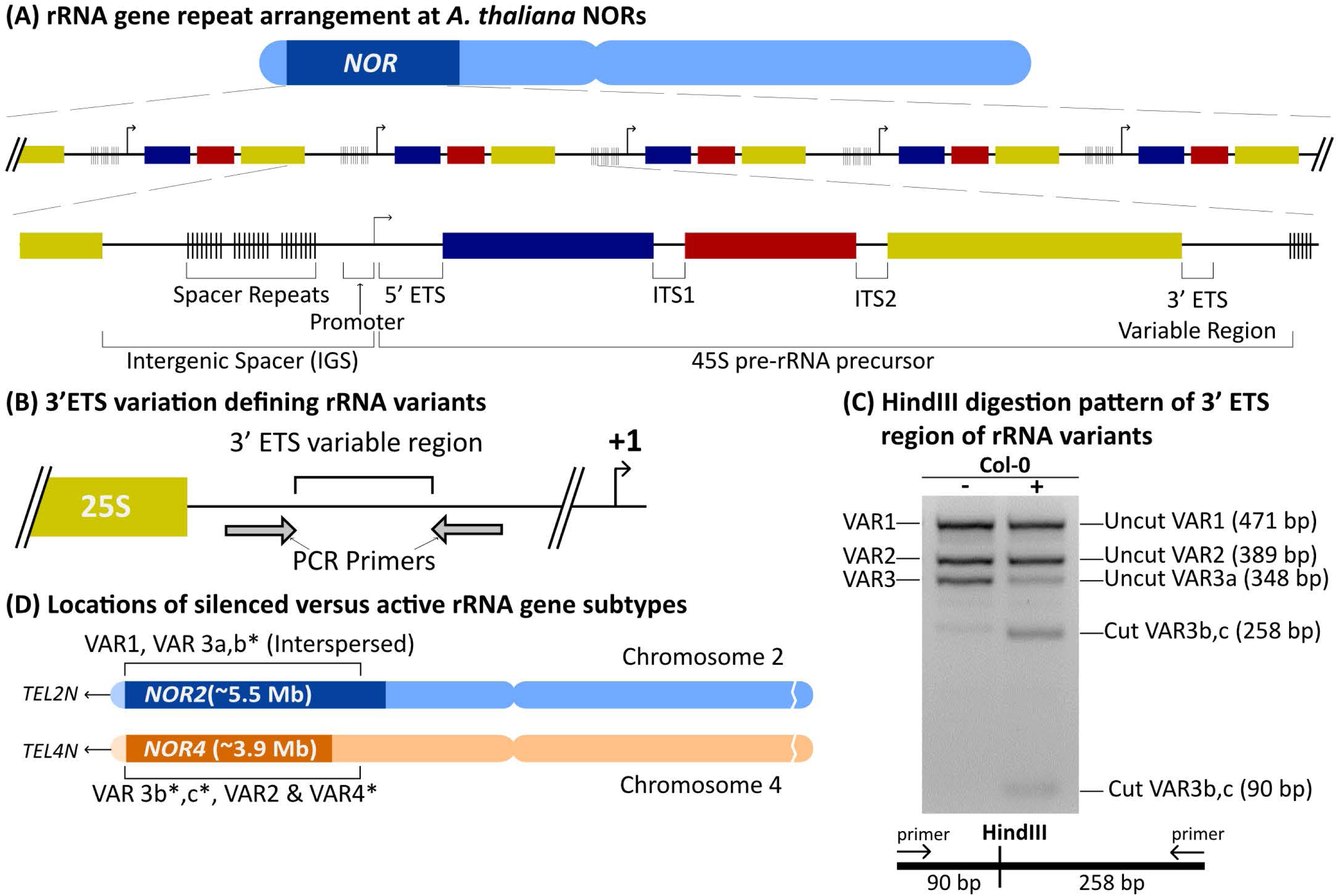
Ribosomal RNA (rRNA) gene organization, subtype variation, localization and their relative arrangement. (A) The diagram shows a generic tandem arrangement of 45S rRNA genes on a NOR. (B) The diagram shows the 3’ ETS region of 45S rRNA genes that exhibit length and sequence polymorphisms among the rRNA variants. (C) The gel image shows HindIll CAPS assay of rRNA variants and the diagram at the bottom shows the position of Hindlll restriction site relative to the binding sites of primers that amplify all four rRNA variants. (D) The diagram shows known locations of the rRNA variants on NOR2 and NOR4 in the Col-0 ecotype of A. thaliana

To determine the relative impact of loss of CG, CHG, and CHH methylation on selective *NOR2* rRNA gene silencing, we analysed rRNA variant expression in mutants defective for DNA methylation, including *ddm1*, *cmt2*, *met1*, *cmt3*, and *drm2* mutants. In these mutants and the WT-Col-0 strain, we performed RT-PCR analysis using RNA isolated from 4-week-old leaf tissues. VAR1 type rRNA genes constitute ∼50% of total 45S rRNA genes, and most of them map to *NOR2* and undergo selective silencing during development (Chandrasekhara et al., 2016, Pontvianne et al., 2010). Our results showed that among the DNA methylation mutants, VAR1 expression levels were found to be higher in *cmt2, ddm1,* and *met1* mutants than the other mutants (Figure 2, Figure S1). In contrast, the *cmt3* and *drm2* mutants have displayed modest levels of VAR1 expression (Figure 2, Figure S1).

**Figure 2.**
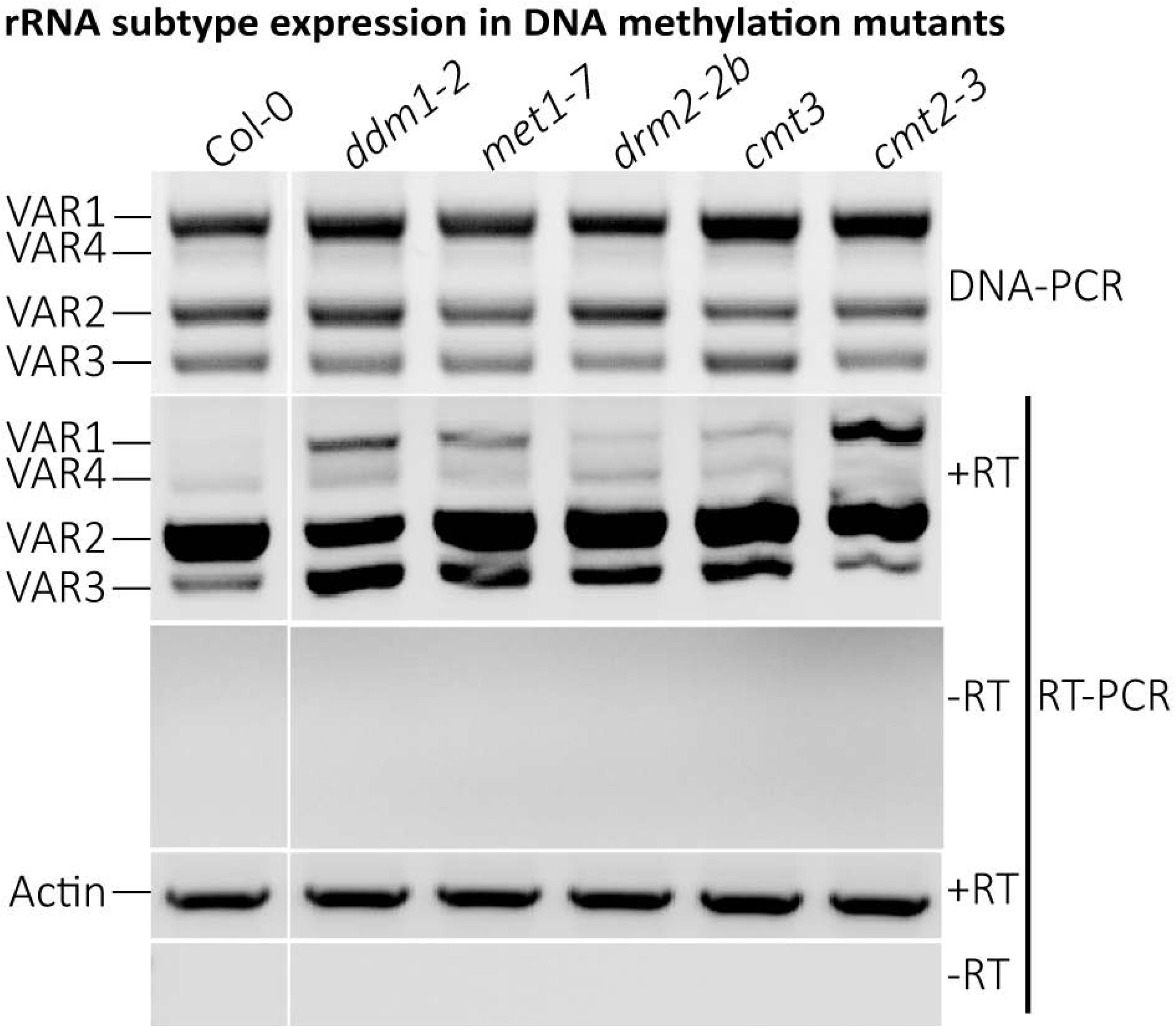
Disruption of *NOR2* gene silencing to varying degrees in DNA methylation mutants. The gel images show DNA-PCR data for rRNA gene subtypes and RT-PCR expression data for rRNA gene subtypes and actin. The gel images were spliced to remove the second lane.

### *NOR2* rRNA genes are hypermethylated while *NOR4* rRNA genes are hypomethylated in Col-0

We performed methylation analysis of rRNA variants using HpaII-digested genomic DNA (gDNA) from Col-0 and *met1* mutants (Figure 3). Because the rRNA variant-defining 3’ ETS region lacks HpaII sites, HpaII digestion of gDNA does not impact PCR amplification of this region. Therefore, we chose to separate HpaII-digested and HpaII-undigested fractions of gDNA by resolving them on an agarose gel. The fragments that remained at the top of the gel (>20 kb) constituted the HpaII-undigested fractions, while those resolved at the 0.6-1.0 kb range constituted the HpaII-digested fractions (Figure 3A, left panel). Both the digested and the undigested fractions were gel-eluted, followed by PCR amplification of rRNA variants. Through this analysis, we identified the rRNA variants that were enriched in either HpaII-digested or undigested fractions. The PCR data revealed VAR2 genes from *NOR4* to be the most predominant in Col-0 among the HpaII-digested products, indicating relative hypomethylation of *NOR4* genes compared to *NOR2*-localized VAR1 genes (Figure 3A, lanes 1 and 2, right panel). However, in *met1* mutants, used as a control, all rRNA variants were amplified proportionate to their composition due to loss of cytosine methylation in all of them (Figure 3A, lanes 3 and 4, right panel,).

**Figure 3.**
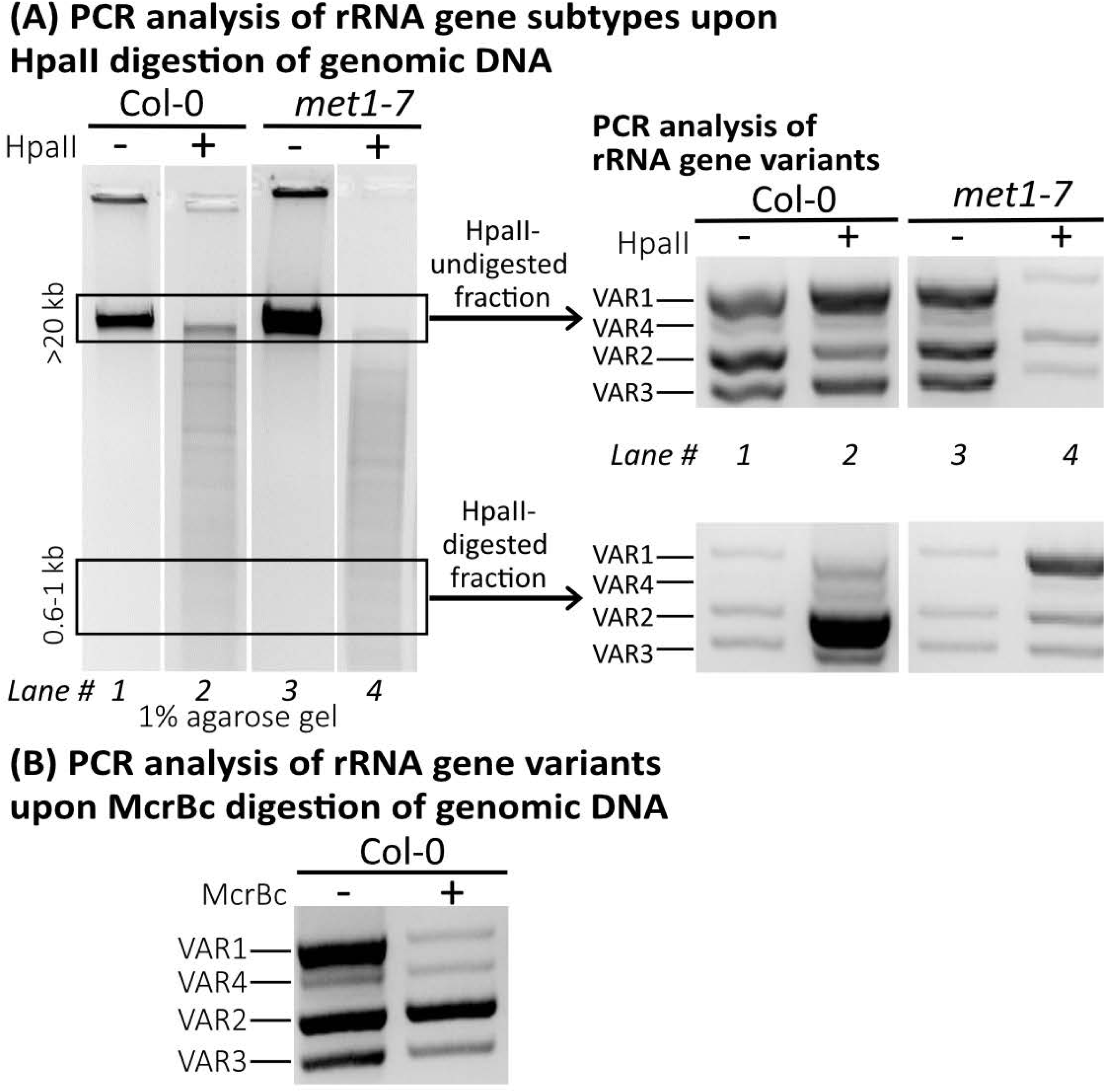
*NOR2* genes are hypermethylated and *NOR4* genes are hypomethylated. (A) Gel images on the left show resolution of Hpall- or buffer-treated Col-0 and *metl-7* DNA samples and the gel images on the right show PCR amplification ofrRNA gene subtypes. For these PCR reactions, gel-eluted DNA samples representing the Hpall-undigested (≥ 20 kb) and the Hpall-digested (0.6-1.0 kb) fractions were used. (B) The gel image shows rRNA gene subtype PCR products obtained from McrBc-digested or no enzyme-treated Col-0 DNA.

In another experiment, we subjected gDNA to digestion by McrBC, an enzyme that preferentially digests DNA if its recognition site (R^m^C(N_40-3000_)R^m^C) consists of methylated cytosines. McrBC digestion of gDNA followed by PCR amplification of rRNA variants revealed that VAR1 was digested the most and VAR2 the least (Figure 3B), indicating that *NOR2* genes are hypermethylated in contrast to *NOR4* genes, consistent with recent findings in *Arabidopsis* which employed sequencing-based methylation analysis of rRNA variants (Fultz et al., 2023, Mo et al., 2023).

### *NOR2* rRNA genes are hypermethylated, while *NOR4* rRNA genes are hypomethylated in the introgression line Col^Sf-^*^NOR4^*

In a previous study, we demonstrated that an introgression line named Col^Sf-NOR4^ comprises *NOR2* from Col-0 (containing VAR1 and VAR3 genes) and *NOR4* from the Sf-2 ecotype (Chandrasekhara et al., 2016) (Figure 4A). Sf-2 has only VAR1 type of rRNA genes, but they can be differentiated from VAR1 genes of Col-0 by HindIII digestion. Sf-2 VAR1 genes have a HindIII site in their 3’ ETS region, while Col-0 VAR1 genes lack it (Chandrasekhara et al., 2016). In Col^Sf-NOR4^, Sf-2 *NOR4* genes are active, though they are similar in sequence to the silenced Col-0 *NOR2* (VAR1) genes (Chandrasekhara et al., 2016). However, Col-0 *NOR2* genes are still selectively silenced in Col^Sf-NOR4^ (Chandrasekhara et al., 2016). Therefore, we analyzed this introgression line to determine if VAR1 type of genes residing in *NOR2* and *NOR4* of Col^Sf-NOR4^ are differentially methylated or not, using similar methylation assays as described above. In one experiment, gDNA of Col^Sf-NOR4^ was subjected to HpaII digestion, followed by gel elutions of the undigested and the digested fractions. The gel-eluted DNA samples were then subjected to PCR amplification of rRNA variants followed by HindIII digestion. In control samples (no HpaII), HindIII digestion of PCR-amplified rRNA variants resulted in the expected products, which include digested Sf2-VAR1 genes, undigested Col-0 VAR1 genes, and partially digested Col-0 VAR3 genes (Figure 4B, lanes 1 and 2, top right gel image, and Figure 4C) (Chandrasekhara et al., 2016). However, HindIII digestion of the PCR products obtained from the gel-purified HpaII-digested fraction as the template revealed that Col-0 VAR1 genes predominantly constitute the HpaII-undigested fraction (Figure 4B, lanes 3 and 4, top right gel image, and Figure 4C). On the other hand, we observed a significant increase in the Sf-2 VAR1 genes in the HpaII-digested fraction (Figure 4B, lanes 3 and 4, bottom right gel image, and Figure 4C) compared to the undigested fraction (Figure 4B, lanes 3 and 4, top right gel image, and Figure 4C). These data indicate hypermethylation of Col-0 *NOR2* genes and hypomethylation of Sf-2 *NOR4* genes, despite the presence of the VAR1 type of genes on both the NORs in the Col^Sf-NOR4^ introgression line.

**Figure 4.**
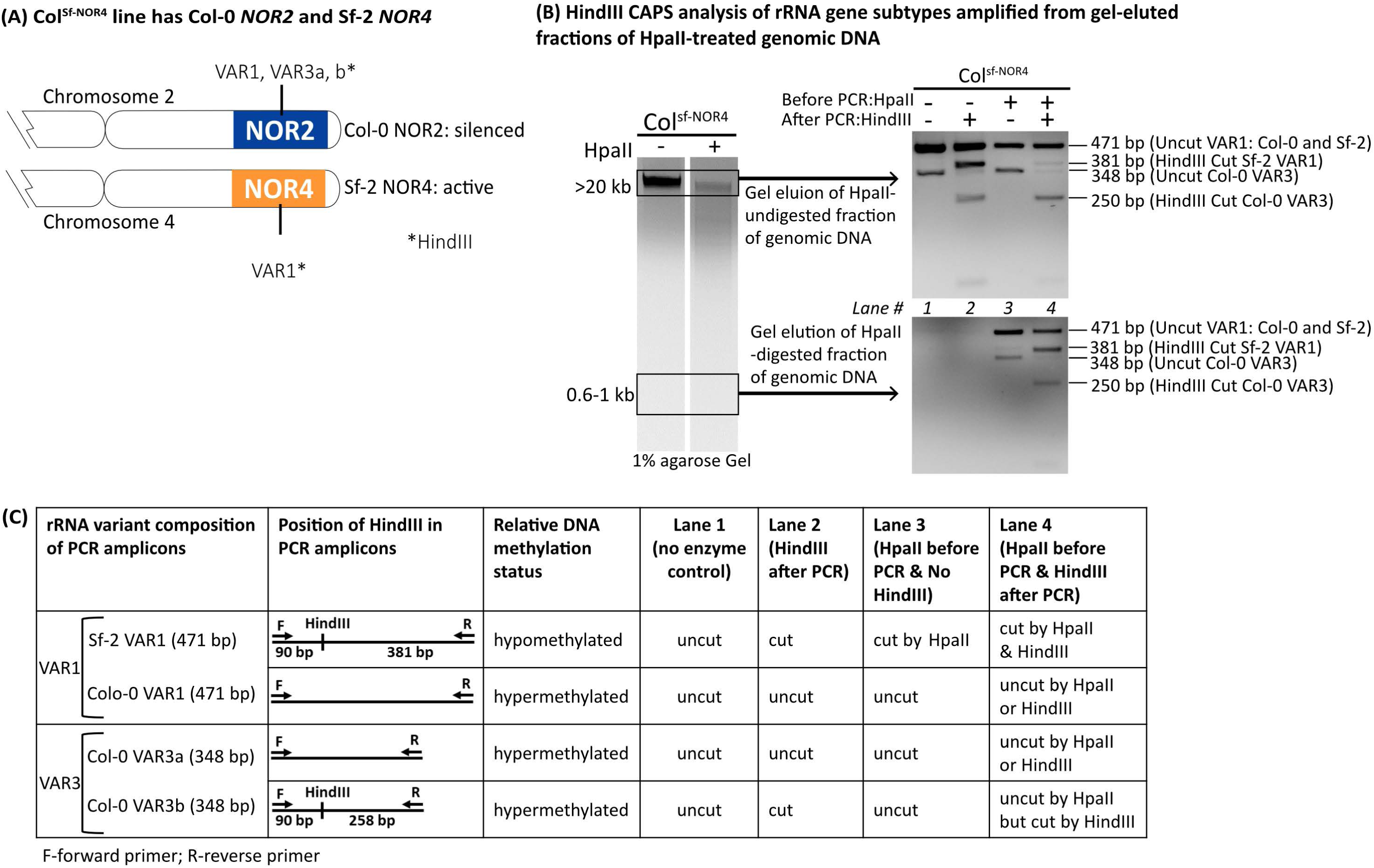
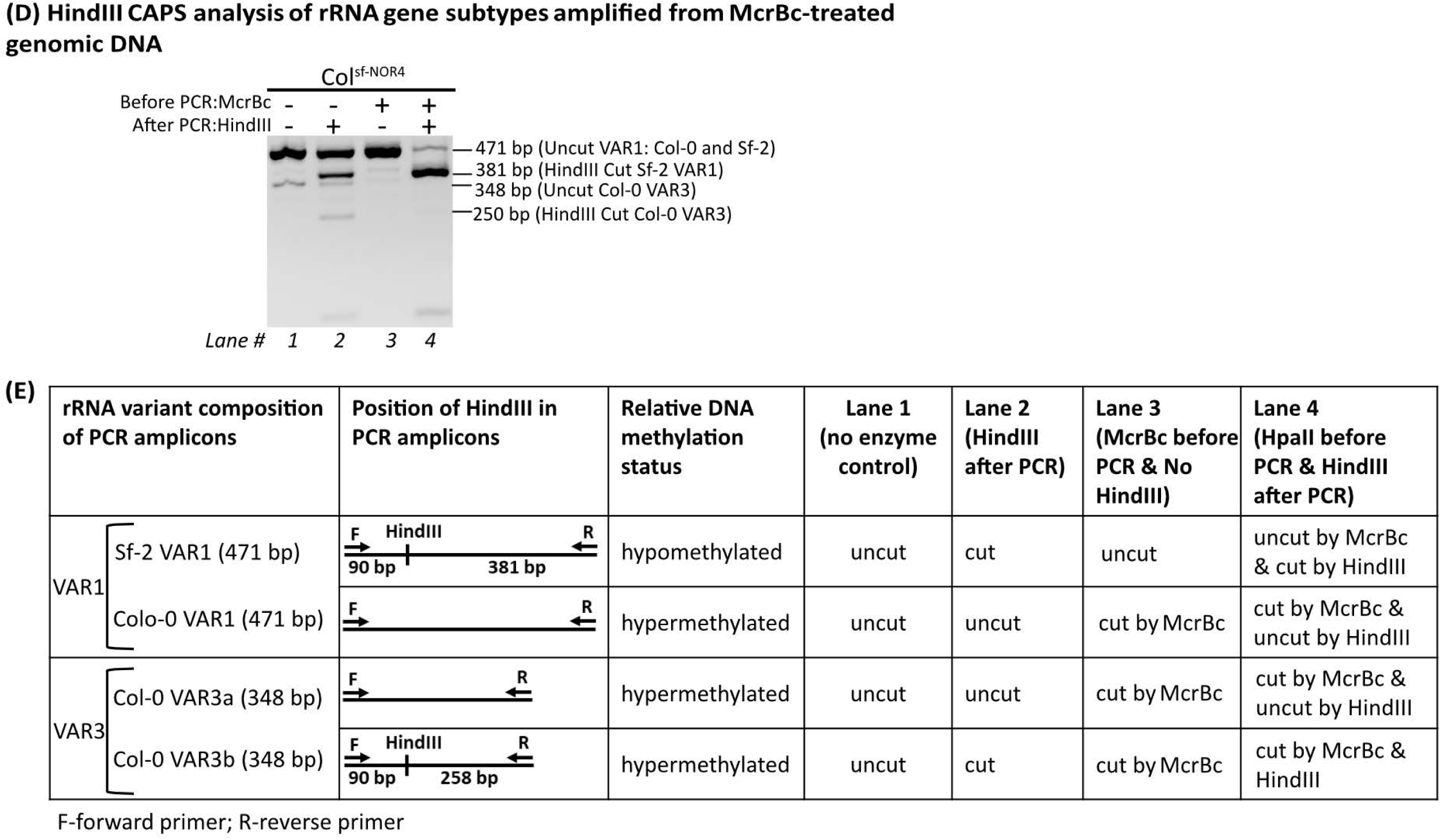
Col-0 *NOR2* genes are hypermethylated and Sf-2 *NOR4* genes are hypomethylated in the Col^st-NDR4^ introgression line. (A) Diagram showing rRNA subtype composition of the Arabidopsis introgression line called CoI^sf-NOR4^ in which Sf-2 *NOR4* and adjacent sequences were introgressed into the Col-0 genetic background. (B) The agarose gel images on the left show resolution of Hpall- or buffer-treated Col^sf-NOR4^ gDNA samples, and the gel images on the right show PCR amplification of rRNA variants with or without Hindlll digestion. For these PCR reactions, gel-eluted DNA samples representing the Hpall-undigested (;:,, 20 kb) and the Hpall-digested (0.6-1.0 kb) fractions were used. (C) Summay of results shown in (B). (D) The Gel image shows PCR amplifica-tion of rRNA variants obtained from McrBc- or buffer-treated co1^sf-NOR4^ gDNA with or without HindiII digestion. (E) Summary of result shown in (D).

In another experiment, gDNA of Col^Sf-NOR4^ was first digested with McrBC, followed by PCR amplification of rRNA variants. We then digested these PCR products with HindIII and resolved them using regular gel electrophoresis. HindIII-digested control PCR products (no McrBC digestion) behaved as expected based on the presence or the absence of HindIII sites (Figure 4D-E, lanes 1 and 2). However, the McrBC digestion patterns of the rRNA variants (Figure 4D-E) were the opposite of the experimental results involving HpaII digestion (compare to Figure 4B-C). Predominantly, the Sf-2 VAR1 genes constituted the McrBC-undigested fraction, while the Col-0 VAR1 genes constituted the McrBC-digested fraction (Figure 4D-E, lanes 3 and 4). We also observed complete digestion of the Col-0 VAR3 (from *NOR2*) genes by McrBC (Figure 4D, lane 3). These data again demonstrate that *NOR4* genes are hypomethylated and *NOR2* genes are hypermethylated in Col^Sf-NOR4^.

In Figures 3 and 4, resistance to HpaII digestion indicates the presence of CG and CHG methylation, while digestion by McrBC indicates the prevalence of CG, CHG, and CHH methylation. Based on these data, we conclude that *NOR2* genes are hypermethylated compared to *NOR4* genes in all cytosine contexts, and this status do not change irrespective of the type of rRNA variants present on *NOR2* or *NOR4*.

### The rRNA gene methylation heat maps reveal a critical role for CMT2-mediated CHH methylation in rRNA gene silencing

Using published whole-genome bisulfite sequencing data (Stroud et al., 2013), we generated heatmaps of cytosine methylation for a 45S rRNA reference gene sequence (Data File S1) using Methylpy (Schultz et al., 2015). DDM1 is required for cytosine methylation in all sequence contexts predominantly in heterochromatin than euchromatin (Jeddeloh et al., 1999, Lippman et al., 2004, Vongs et al., 1993, Zemach et al., 2013). *NOR2* primarily exists as heterochromatin while *NOR4* exists as euchromatin (Figure 3)(Fultz et al., 2023, Mo et al., 2023), which correlates with their differential rRNA gene expression (active *NOR4* vs inactive *NOR2*)(Fultz et al., 2023, Mohannath et al., 2016, Chandrasekhara et al., 2016). Therefore, we presume that the loss of cytosine methylation shown in methylation heat maps for all the mutants (Figure 5) largely reflects losses from *NOR2* than *NOR4*.

**Figure 5:**
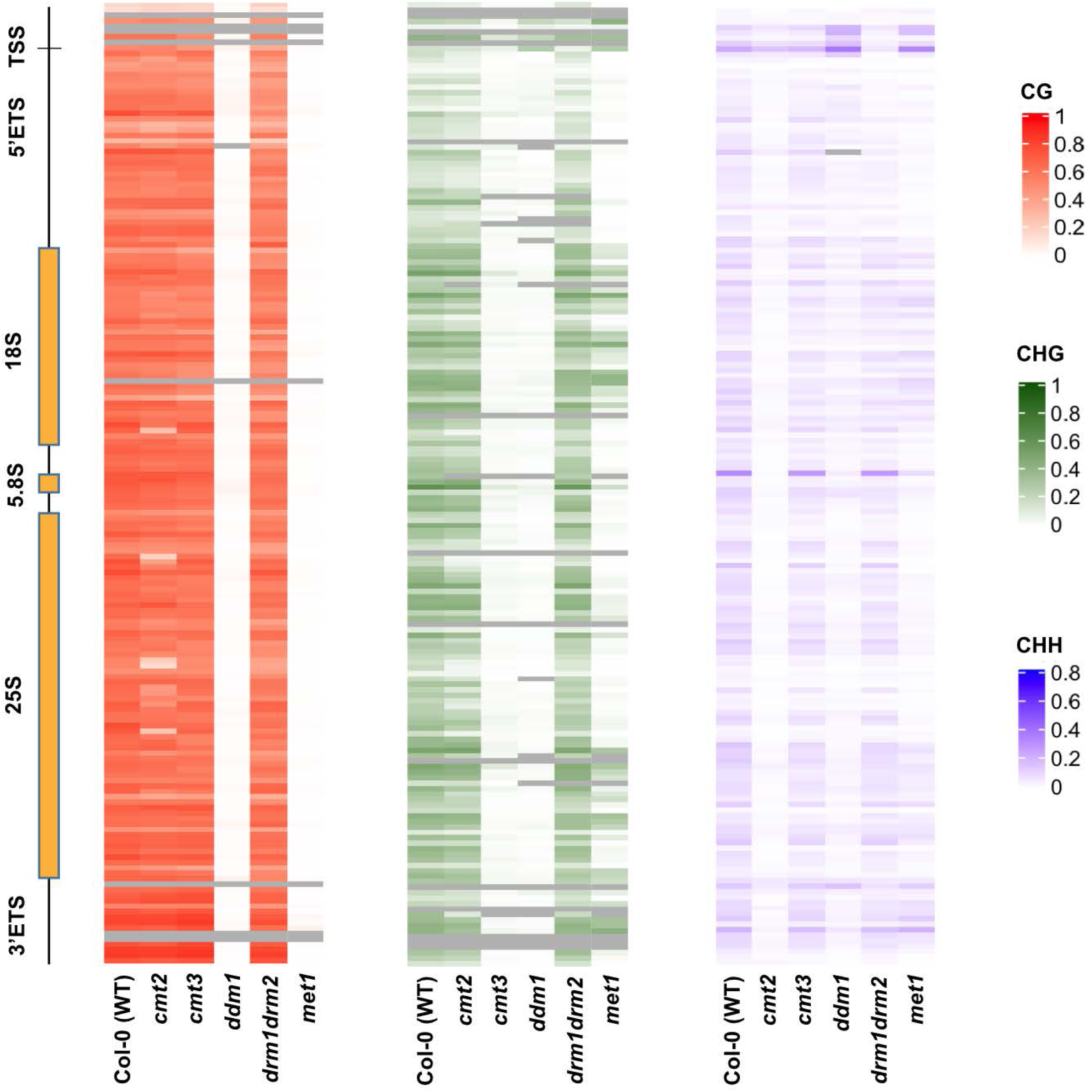
Methylation state of rDNA in CG, CHG, and CHH contexts. Whole genome bisulfite sequencing data from various DNA methylation mutants were analyzed using Methylpy, and the overall cytosine methylation state of rDNA is shown as heatmaps in all three cytosine contexts. The Y-axis represents a single rDNA unit starting with a promoter and ending with 3’ ETS. Each line on the heatmap represents the methylation state of the cytosines in a given methylation context in 50 nucleotide bins. CG is shown in red, CHG in green, and CHH in blue. Grey indicates that no methylation sites were present in that bin for a given context. TSS= Transcription Start Site

The rDNA cytosine methylation heatmaps show that the entire rRNA gene sequences are methylated in all sequence contexts (Figure 5, see Col-0 column in CG, CHG, and CHH tracks), consistent with recent findings (Fultz et al., 2023). Among the mutants, the highest overall loss of CHH methylation was observed in *cmt2*, followed by *ddm1* and *met1* mutants (Figure 5). Indeed, these mutants also show higher *NOR2* gene silencing disruption than the other mutants (Figure 2, Figure S1). Interestingly, in *drm1drm2* mutants, although the overall CHH methylation loss is significantly lesser than that of *cmt2* mutants, the losses are comparable around the gene promoter (GP) and the upstream region or are even slightly higher in *drm2* mutants (Figure 5). These results indicate two things; one, RdDM-independent CMT2, primarily methylates CHH sites throughout the gene sequences, while RdDM-dependent DRM primarily methylates CHH sites present in the GP and its upstream sequences. Second, despite the losses of CHH methylation around the promoter region being comparable between *cmt2* and *drm2* mutants (Figure 5), the disruption of rRNA gene silencing is significantly more in *cmt2* than in *drm2* mutants (Figure 2), suggesting a role for gene body CHH methylation in rRNA gene silencing. Recent studies, including one on DDM1, demonstrate that gene body methylation also play a role in gene silencing (Lee et al., 2023a, Shahzad et al., 2025, Williams et al., 2023). Further, the CG methylation levels in *cmt2* mutants are comparable to those of WT-Col-0 (Figure 5), yet the loss of *NOR2* gene silencing is among the highest in *cmt2* mutants (Figure 2, Figure S1), indicating that CHH methylation loss is sufficient for releasing rRNA gene silencing. As a corollary, the presence of CG methylation is not sufficient for rRNA gene silencing.

The *met1* and *ddm1* mutants showed near complete loss of CG methylation at rDNA loci, while other mutants hardly lost any CG methylation (Figure 5). Also, the *met1* mutants showed some loss of CHH and CHG methylation at few sites within rRNA gene body sequences but gained some CHH methylation in the gene promoter region, not in the gene body (Figure 5). Given these data, we attribute the significant disruption of *NOR2* gene silencing observed in *met1* mutants (Figure 2) to the near-complete loss of CG methylation in them (Figure 5), without ruling out a minor contribution from the CHH methylation loss observed in a small portion the gene body. CHH hypermethylation around the promoter region has also been observed In *hda6* mutants which display a loss of CG and CHG methylation and disruption of *NOR2* gene silencing (Chandrasekhara et al., 2016, Earley et al., 2010, Mohannath et al., 2016). Consistent with our conclusion that both CG and CHH methylation are required for rRNA gene silencing, we hypothesize that the disruption of *NOR2* gene silencing observed in *hda6* mutants is at least partly due to their CG methylation loss (Earley et al., 2010). Moreover, another study indicates that HDA6 could affect gene silencing without significantly altering DNA methylation (Hristova et al., 2015).

Next, given the modest release of *NOR2* rRNA gene silencing in *cmt3* mutants despite near complete loss of CHG methylation in them (Figures 2 and 5), we conclude that the CHG methylation has a minimal role in rRNA gene silencing. Based on the results discussed so far, we propose that combination of MET1-mediated CG and CMT-mediated, RdDM-independent CHH methylation is required for selective rRNA gene silencing. Comparable losses of *NOR2* gene silencing disruption in *ddm1* mutants, which display near complete loss of cytosine methylation in all sequence contexts, and *cmt2* mutants, which display a loss of most of CHH methylation (Figures 2 and 5), indicate that MET1-mediated CG and CMT2-mediated CHH methylation act in a complementary manner, rather than synergistically, in rRNA gene silencing.

Another observation of some interest is that lower levels of CHG and CHH methylation losses at rDNA loci in *met1* mutants are not uniform, rather they occur at the same two sites for the CHG and CHH methylation losses (Figure 5). One site encompasses the 5’ETS region, while the other starts near the end of 18S and ends near the middle of 25S (Figure 5, compare CHG and CHH tracks of *met1* mutants). Interestingly, in the latter region, *cmt2* mutants also show some loss of CG methylation (Figure 5, compare CHH track of *met1* mutants and CG track of *cmt2* mutants). These results suggest as if some specific sites in rRNA gene body require both CMT2 and MET1 for efficient CG and CHH methylation, but we are not aware of any such existing links between these two methyltransferases.

The data depicted in the methylation heat maps (Figure 5) depicting varying losses of CG, CHG, and CHH methylation at rDNA loci and disruption of *NOR2* gene silencing in *met1*, *cmt3*, and *drm2* mutants are consistent with findings from previous studies (Cokus et al., 2008, Pontvianne et al., 2013). But notably, *ddm1* and *cmt2* mutants were not included in these studies. Studying these mutants proved crucial in establishing a critical role for CMT2-mediated CHH methylation, in combination with CG methylation, in rRNA gene silencing.

### Cytosine occurrence in rRNA genes is the highest in the CHH context followed by CG and the lowest in the CHG context

We wondered why CHH methylation loss significantly impacts *NOR2* rRNA gene silencing, despite the *NOR2* genes showing relatively lower levels of CHH methylation (Fultz et al., 2023, Mo et al., 2023). To address this question, we analyzed the distribution of cytosines in CG, CHG, and CHH sequence contexts across all the 45S rRNA gene sequences, obtained from a recent study (Fultz et al., 2023). From these sequences, we depict 2 reference sequences, which commonly represent *NOR2* and/or *NOR4* (Figure 4A and Data File S2). We analyzed the gene sequences in five categories: gene promoter (GP), spacer promoter (SP), GP upstream sequence, Sal clusters and other sequences of intergenic space (IGS), and the gene body, which includes the entire transcribed (pre-45S rRNA) sequence. In *A. thaliana*, the −55 to +6 region with respect to the transcription start site (+1) in the 45S rRNA genes constitutes the minimal rRNA gene promoter (hereafter referred to as the gene promoter) (Doelling et al., 1993, Doelling and Pikaard, 1995), which occurs only once per gene repeat (Figure 4A). The GP is 100% conserved in 96% of the gene repeats, while the remainder of the repeats carries one to a few mismatches (Fultz et al., 2023). The GP has eight cytosines, but strikingly, none of them exist in the CG context. Six (75%) exist in the CHH context, and the remaining two (25%) exist in the CHG context (Figure 4A).

Similarly, we analysed the predicted GP regions in the recently assembled tomato *NOR2*, constituting ∼1500 45S rRNA genes (Shirasawa and Ariizumi, 2024), and the previously published tomato 45S rRNA gene promoter sequence (Doelling and Pikaard, 1995). 98% of the gene repeats show 100% homology to the predicted GP sequence (−55 to +6) and the remaining repeats showed one to few mismatches (Shirasawa and Ariizumi, 2024). This predicted GP has 12 cytosines, of which 10 (∼83%) exist in the CHH context, 2 (∼17%) in the CHG context, and none in the CG context (Figures 4B-C and S2, Data File S3) (Doelling and Pikaard, 1995, Shirasawa and Ariizumi, 2023, Shirasawa and Ariizumi, 2024). Similar cytosine distributions were found in the predicted GPs of other plant species (Figure 4B-C) (Doelling and Pikaard, 1995). Further, we analyzed ∼270 bp immediate upstream region of the GP, in which, out of 45-46 cytosines, 4 each (9% each) exist in CG and CHG contexts, while 35-36 (81-82%) exist in the CHH context (Figure 4A). Likewise, in the immediate upstream region of the predicted GP in tomato 45S rRNA gene sequence up to 413 bp, cytosine distribution is largely skewed towards CHH sites. Out of 43 cytosines, we found 10 (23%) in CG, 3 (7%) in CHG, and 30 (70%) in CHH context (Figure S2).

Next, we tabulated cytosine distribution in the Arabidopsis spacer promoters (SPs), whose number varies from 0 to 4, with 1 SP being the most common in both *NOR2* and *NOR4* (Figure 4A) (Fultz et al., 2023). The SPs share homology with the GP sequence and can program low-level transcription by RNA polymerase I (Doelling et al., 1993). Each SP has a total of 14 cytosines, of which 11 (∼79%) are in the CHH context, 2 (∼14%) are in the CHG context, and 1 (∼7%) in the CG context (Figure 4A, Data File S2). Further, we calculated cytosine distribution in other regions of IGS, including the Sal clusters, which are made of 20-21 bp repeats containing a SalI restriction site, and other interspersed sequences (Fultz et al., 2023). Interestingly, in this region, we found predominant occurrence of CG (52-53 %) over CHG (25-26 %) and CHG (21-22 %) sites, even among IGS regions varying in length from ∼350 to 465 (Figures 4A, Data File S2). If we consider the total cytosines across the entire 45S rRNA gene sequence, the frequency of CHH sites (45-46%) ranks first while the frequencies of CG (35%) and CHG sites (19-20%) rank second and third, respectively (Figure 4A).

Similar analysis of tomato IGS sequences did not yield any IGS elements containing Sal repeats or sequence elements which shared homology with the predicted GP. In the predicted IGS of tomato, although we found that the number of CHH sites(139/256) is still the highest among three contexts, the number of CG sites (99/256) increase substantially in this region (Figure S2). The number of CHG sites (18) is the lowest (Figure S2). Overall cytosine distribution pattern in tomato 45S rRNA gene sequences mirror that of Arabidopsis, with number of CHG sites > CG sites > CHG sites (Figure S2).

Recent studies have shown that the gene body methylation also plays a role in gene regulation (Lee et al., 2023a, Shahzad et al., 2025, Williams et al., 2023), and in Arabidopsis, the entire rRNA gene sequences are heavily methylated (Figure 2) (Fultz et al., 2023). Therefore, we extended cytosine distribution analysis to rRNA gene bodies, wherein we found the occurrence of more CHH sites than CG and CHG, both in *Arabidopsis* (49% CHH > 32% CG > 19% CHG) and tomato (46% CHH > 35% CG > 18% CHG) (Figures 4A and S2, Data Files S2-S3). Similar trends were observed when we considered the entire 45S rRNA gene sequences (*Arabidopsis*: 46% CHH > 35% CG > 20% CHG, tomato: 48% CHH > 35% CG > 17% CHG) (Figures 4A and S2, Data Files S2-S3). Collectively, these findings provide a putative mechanistic basis for the maximum disruption of *NOR2* rRNA gene silencing in mutants that displayed higher loss of CHH (*cmt2, ddm1, and met1*) or CG methylation (*met1* and *ddm1*) than the other mutants. Importantly, these results define a novel critical role for CHH methylation in regulating 45S rRNA genes in plants.

In contrast, when we analyzed human rDNA promoter sequences (Zhou et al., 2016), we found that, out of 65 cytosines, 26 are in CG (∼40%; the highest among the three), 19 in CHG (∼29%), and 20 in CHH (∼31%) contexts (Figure S3, columns 2-3, Data File S4). Even when we included an extended upstream sequence as part of the rDNA promoter, according to another study (Uemura et al., 2012), the cytosine distribution trend remained the same (Figure S3, columns 4-5, Data File S4). Similarly, we found the predominant occurrence of CG sites in the gene bodies (Figure S3, the last two columns, Data File S4). These results are consistent with the mammalian rRNA gene regulation, which is almost entirely dictated by CG methylation (Bernstein et al., 2007, Priyadarshini et al., 2023, Zhou et al., 2016). These findings also correlate with the evolutionary loss of chromomethylases (CMTs) in mammals. CMTs are plant-specific, which primarily methylate CHG and CHH sites (Bewick et al., 2017). Collectively, these findings indicate the presence of evolutionarily divergent mechanisms of rRNA gene silencing in plants and mammals.

## Discussion

The study primarily provides two new insights. First, RdDM-independent CMT2, mediates majority of the CHH methylation at rDNA loci (Figure 5). Second, the loss of CMT2-mediated CHH methylation is sufficient for disrupting selective *NOR2* gene silencing (Figures 2 and 5). A recent study in a different plant species highlights a role of non-CG methylation in plant rRNA gene silencing (Matyášek et al., 2024), but it did not include CMT2. Our data also capture findings from a previous study in underscoring a critical role played by CG methylation in rRNA gene silencing (Figures 2, 3 and 5). Based on our collective findings, we propose a model stating that a combination of CG methylation and CMT-mediated RdDM-independent CHH methylation is required for rRNA gene silencing (Figure 7). Losing any one of them is sufficient for releasing the rRNA gene silencing. In *ddm1* mutants, CG and CHH methylation are nearly wiped out, their NOR2-residing VAR1 expression levels are comparable to that of *cmt2* mutants, in which only CHH methylation is nearly wiped out (Figures 2 and 5). These data prompted us to further proposed that MET1-mediated CG and CMT2-mediated CHH methylation act complementarily, rather than additively, in silencing rRNA genes.

Based on the most abundant CG methylation found at NOR loci (Cokus et al., 2008, Fultz et al., 2023, Mérai et al., 2014, Pontvianne et al., 2013), it is fairly understandable mechanistically how CG methylation could play a role in rRNA gene silencing. However, relative methylation levels of CHH being the lowest at rDNA loci, it has not been obvious how CHH methylation loss would disrupt rRNA gene silencing. To address this gap, we asked an important question-cytosine methylation of which regions play a role in rRNA gene silencing? To address this question, we analyzed cytosine distribution among the three sequence contexts (CG, CHG, and CHH) in regulatory and gene bodies of the rRNA genes. In the characterized regulatory elements constituting GP and SP, CHH site occurrence (17/22) is overwhelmingly high compared to either CG (1/22) or CHG (4/22) and the trend remained the same even when we analyzed ∼270 bp upstream of GP region or the gene body (Figure 6A). Likewise, analysis of comparable regions from tomato rRNA sequences also yielded the similar cytosine distribution heavily in favour of CHH over CG or CHG sites (Figure S2). These findings provide a potential mechanistic basis to explain how CHH methylation could play a critical role in rRNA gene silencing. When the number of CHH sites are overwhelmingly high in a region of regulatory importance, low levels of CHH methylation can be compensated by higher number of CHH sites. For instance, in the GP and SP regions combined, CHH site frequency is 17- and 4.3-fold higher than CG and CHG sites, respectively (Figure 6A). However, Sal-repeats-containing IGS (minus SP) region, is highly CG-rich (CG- 189 sites, CHG- 91 sites, and CHH- 75 sites, when we considered the most commonly occurring 2-Sal cluster scenario) (Figure 6A), which explains why CG methylation loss significantly disrupts rRNA gene silencing (Figures 2, 5 and 6). In the predicted comparable sequences of tomato NORs, we could not find Sal repeats and in this region, CHH occurrence still turned out to be the highest (CG- 109 sites, CHG- 21 sites, and CHH-167 sites) (Figure S2), suggesting predominant occurrence of CG and CHH sites in the IGS and promoter regions, although in varying proportions.

**Figure 6.**
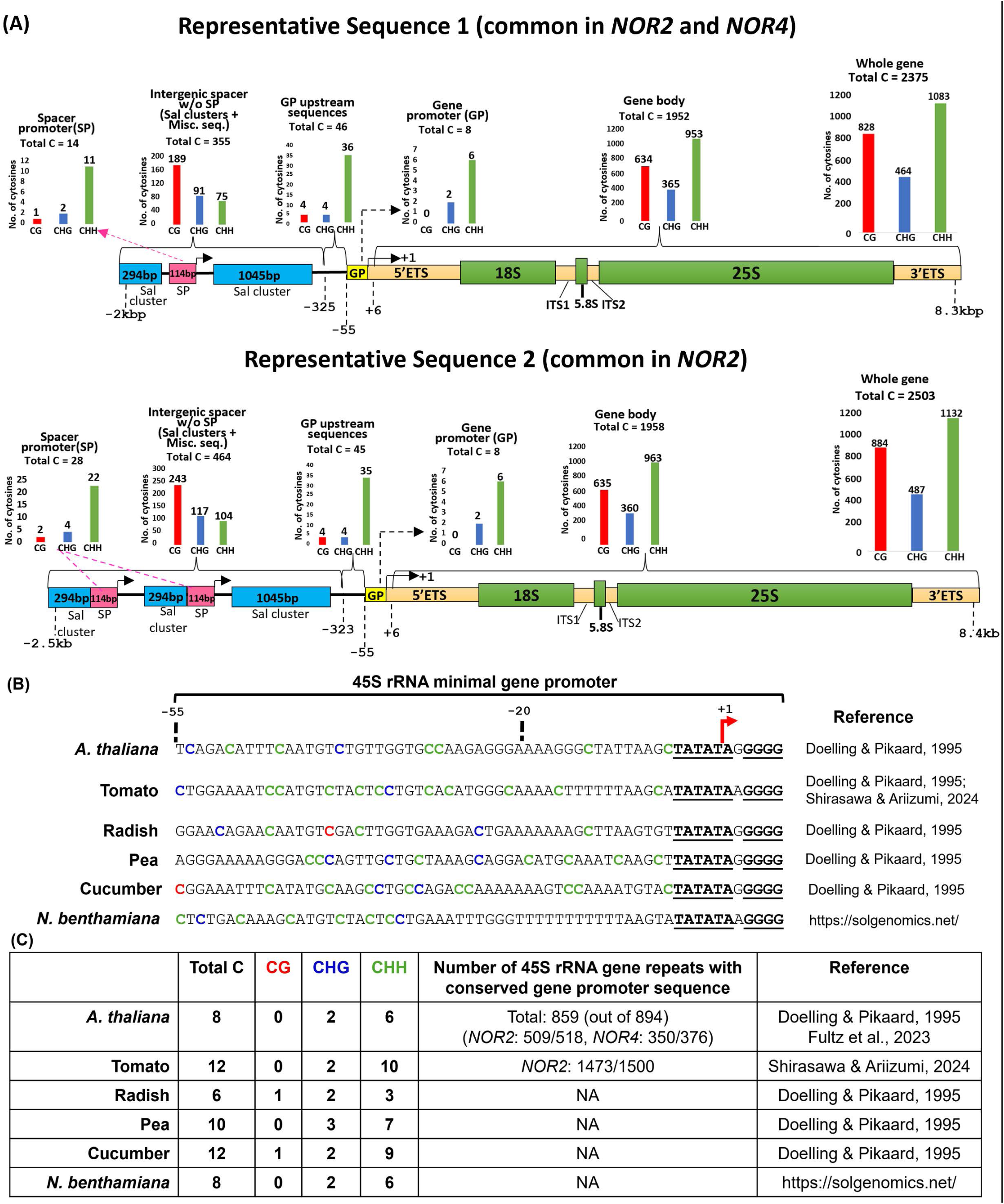
Within the 45S rRNA genes, cytosine occurrence is the highest in CHH context, followed by CG, and the lowest in CHG context. **(A)** A diagram provides cytosine distribution within different regions of 45S rRNA gene sequences. Based on the NOR sequences from Fultz et al., 2023, two reference sequences are chosen for representation based on their common occurrence in NORs **(B)** Sequences show relative distribution of cytosines among different cytosine contexts (CG, CHG, and CHH) in the minimal rRNA gene promoter (GP) regions of different plant species. **(C)** A table shows the number of cytosines in different sequence contexts across different plant species. For *A. thaliana* and tomato, data were obtained from Fultz et al., 2023 and Shirasawa and Ariizumi, 2024, respectively, while the data for other species were obtained from Doelling & Pikaard, 1995. NA-Entire NOR sequences not available.

**Figure 7.**
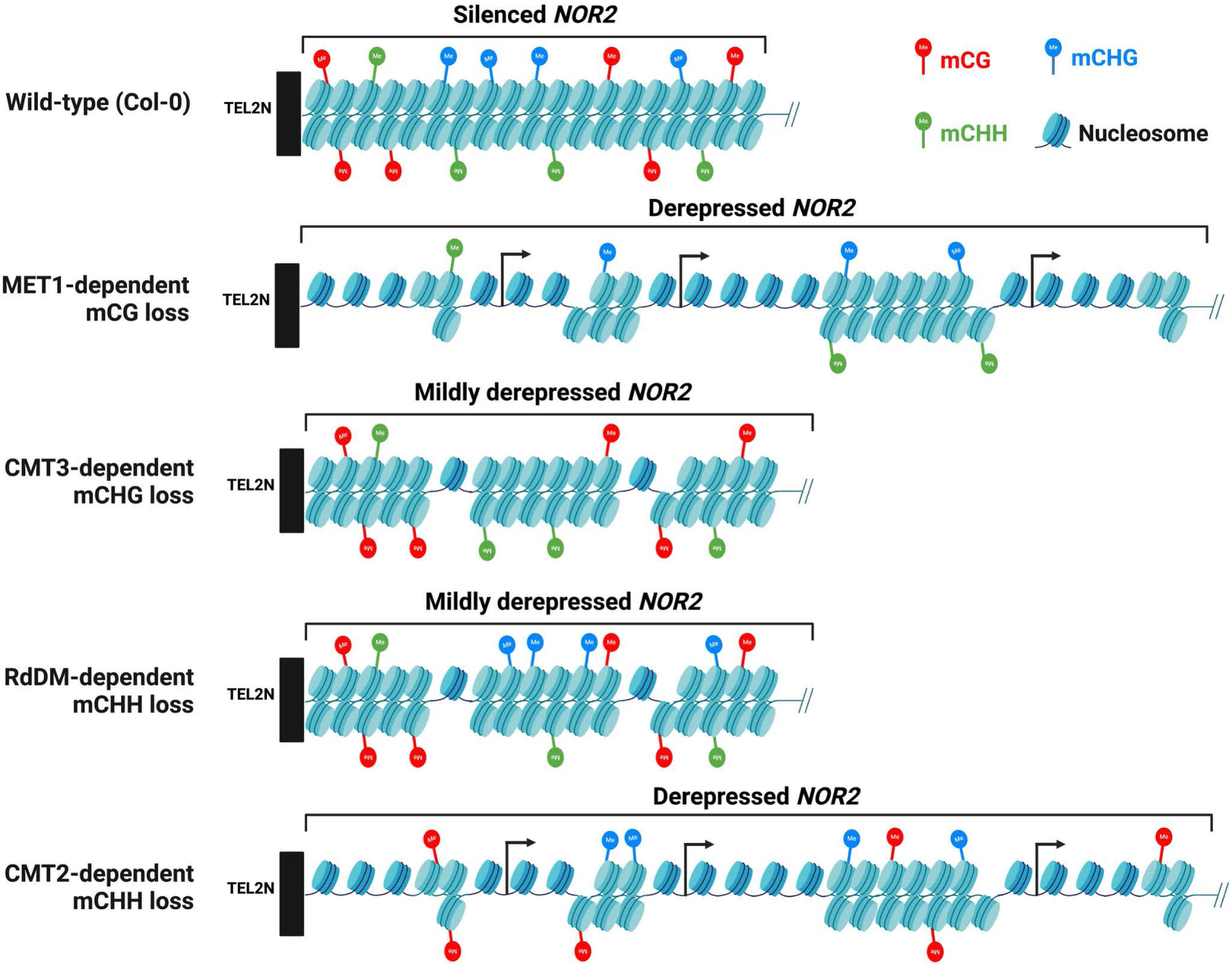
Model depicting a role for CMT2-dependent CHH methylation and other cytosine methylation pathways in rRNA gene silencing in Arabidopsis. In Wild-type Col-0, *NOR2* rRNA genes are methylated in all cytosine contexts. Loss of CMT2-mediated mCHH or METl-mediated mCG results in significant disruption of *NOR2* rRNA gene silencing and likely alleviates chromatin condensation at these loci. However, loss of CMT3-dependent mCHG or RdDM-dependent mCHH has a mild effect on rRNA gene silencing release at *NOR2.* The model takes into account the levels of NOR2 gene silencing disruption (Fig. 2) and the levels of cytosine methylation loss at each context (Fig. 3) for each of the cytosine methylation pathway mutants.

CMT2 and CMT3 methylate DNA through the dual binding of H3K9 di-methylation (H3K9me2)-containing nucleosomes via their bromo adjacent homology (BAH) and Chromo domains (Du et al., 2012). Of note, H3K9 methyltransferases are required for rRNA variant-specific silencing in *A. thaliana* (Pontvianne et al., 2012). However, the loss of CHH methylation at rDNA loci has more impact on *NOR2* derepression than the loss of CHG methylation, presumably due to higher occurrence of cytosines in the CHH context than the CHG context in the rRNA genes (both in the regulatory elements and gene body) (Figures 2, 5, and 6).

DDM1 mediates CHH methylation by CMT2 (Zemach et al., 2013) and therefore it is reasonable to find levels of CHH methylation loss and *NOR2* gene silencing in *ddm1* mutants comparable to that of *cmt2* mutants (Figures 2-3). Recent findings have shown that DDM1 deposits histone variant H2A.W, which marks plant heterochromatin, onto potentially mobile transposable elements (TEs), resulting in their transcriptional gene silencing (Osakabe et al., 2021). We speculate that a similar mechanism, involving DDM1-mediated histone H2A.W deposition, is also involved in *NOR2* rRNA gene silencing.

Our data also indicate a role for gene body methylation in rRNA gene silencing. For example, in *cmt2* and *drm2* mutants, CG methylation is largely intact, little to no change in CHG methylation and comparable loss of CHH methylation in the promoter and the upstream sequences but gene body CHH methylation loss and NOR2 gene silencing disruption are significantly higher in *cmt2* than *drm2* mutants (Figures 2 and 5). These data suggest a role for gene body CHH methylation in rRNA gene silencing. Similar role for gene body CG methylation is highly likely, given its role in the regulation of various genomic loci (Lee et al., 2023a, Shahzad et al., 2025, Williams et al., 2023).

In contrast, rRNA gene silencing is entirely mediated by CG methylation in mammals. For example, human rDNA promoters are mostly methylated in the CG context and their dysregulation has been linked to several types of cancers (Priyadarshini et al., 2023, Bacalini et al., 2014, Ghoshal et al., 2004, Karahan et al., 2015, Raval et al., 2012, Zhou et al., 2016). Moreover, cytosines occur predominantly in the CG context in the human rRNA gene promoters and gene bodies (Figure S3). CHG and CHH methylation are largely absent in mammals, which correlates with their evolutionary loss of CMTs that primarily methylate CHG and CHH (Bernstein et al., 2007, Bewick et al., 2017, Henderson and Jacobsen, 2007). Notably, spacer promoters in animal models (e.g., frog species *Xenopus laevis*) significantly contribute to the enhancement of rRNA gene transcription (De Winter and Moss, 1986). In contrast, spacer promoters have minimal contribution to rRNA gene transcription in plant species (e.g. *Arabidopsis thaliana*) (Doelling et al., 1993). These findings indicate the evolution of divergent rRNA gene regulatory pathways in plants and animals.

The silent *NOR2* genes are hypermethylated in all cytosine contexts (CG, CHG, and CHH) and exist in chromatin with decreased accessibility state than the active *NOR4* rRNA genes, irrespective of the type of rRNA variants on them (Figures 3-4) (Fultz et al., 2023, Mo et al., 2023). Despite the striking similarities in rRNA gene sequences between *NOR2* and *NOR4*, why rRNA genes on *NOR2* are selectively hypermethylated and silenced? It has been speculated that silencing from a stretch of NOR2-flanking transposable elements and transposon remnants that stretch for ∼60 kb before the first protein-coding genes are encountered, could potentially spread to *NOR2* (Chandrasekhara et al., 2016). This ∼60 kb region is heavily cytosine methylated and carries histone post-translational modifications, which are indicative of transcriptionally repressed heterochromatin. In contrast, NOR4 is flanked by a few transposon-related sequences, with the occurrence of active protein-coding genes within ∼3 kb of the NOR4 boundary. However, the question remains to be answered.

## Materials and Methods

### Plant material and growth

*A. thaliana* ecotype Col-0 (stock no. CS22681), and mutants *met1-7* (stock no. SALK_076522), *cmt2-3* (stock no. SALK_012874C), and *cmt3-11t* (stock no. SALK_148381), were obtained from the *Arabidopsis* Biological Resource Center (ABRC; http://abrc.osu.edu). *drm2-2b* (stock no. SAIL_70_E12) and Col^Sf-NOR4^ (Chandrasekhara et al., 2016, Lee and Amasino, 1995) were provided by Andrzej Wierzbicki and Scott Michaels, respectively, and *ddm1-2* was provided by Eric Richards. Plants were grown in soil in a plant growth room under a 16-h light/8-h dark photoperiod cycle with 50–55% humidity and temperature of 23–25°C.

### RNA isolation

Four-week-old leaf tissue of individual plants was frozen in liquid nitrogen and ground into a fine powder. RNeasy Plant Mini Kit (Qiagen) was used to extract total RNA from ground powder. Total RNA was then treated with Turbo DNA-free kit (Invitrogen) to eliminate contaminating DNA.

### PCR and RT-PCR assays

For PCR of genomic DNA to amplify all rRNA variants, PCR amplification was conducted using 22-30 cycles (depending on the source of DNA) of 30 sec at 95°C, 30 sec at 55°C–60°C, and 60 sec at 72°C. For RT–PCR, the reverse transcription reaction was performed using 1 μg of total RNA using SuperScript III reverse transcriptase (Invitrogen). One microliter (∼100 ng) of reverse transcription product was then used for PCR amplification of rRNA variable regions or actin gene control using 30 cycles of 30 sec at 94°C, 30 sec at 55°C, and 60 sec at 72°C. In all the PCR or RT-PCR reactions, primers 5’allvar and 3’allvar were used to amplify rRNA variable regions (all four variant types). PCR and RT–PCR products were resolved by electrophoresis on 2%–2.5% agarose gels in TBE buffer. Sequences of primers used in this study are given in Figure S10. Pixel intensities for bands in Figures 1, S2, 4B, and 5B were calculated using Quantity one software (Bio-Rad).

### Generation of DNA methylation heat maps and rRNA gene cytosine methylation frequencies

Whole genome bisulfite sequencing data (GSE39901) published by Stroud et al. (2013)(Stroud et al., 2013), was used to assess the methylation state of rDNA in various methylation mutants. The sequencing data were analysed using Methylpy (Schultz et al., 2015). The reads were mapped to a converted 45S rDNA reference sequence, allowing a single mismatch. The methylation state of all cytosines in CG, CHG, and CHH contexts was then analyzed and represented as a fraction of methylated cytosines over the total cytosines in each context. Replicates for a given genotype were merged into a single file.

### DNA isolation from leaf tissue and agarose gel pieces (gel elution)

Genomic DNA (gDNA) was isolated from the leaf tissue of 4-week-old plants by using Illustra Nucleon Phytopure extraction kit reagents (GE Healthcare). This gDNA was used for HindIII CAPS assay (Figure S1C), PCR of rRNA variants (Figure 1, topmost panel), methylation analysis of gDNA (Figure 4A left panel, Figure 4B, and Figure 5B, left panel). In some cases, following gDNA PCR or digestion by methylation-sensitive enzymes, the products were resolved on an agarose gel and the DNA fragments were gel-purified using QIAquick gel extraction kit (Qiagen). This gel-purified DNA was subsequently used as a template for downstream PCR amplification of rRNA variants (Figure 4B right panel, Figure S8C) or HindIII CAPS assay (Figure 5B right panel). DNA from the EZ-Tn5 transposase-treated nuclei was isolated using the cetyl trimethyl ammonium bromide (CTAB) method.

The sequences of primers and oligos used in this study are listed in Figure S4.

## Acknowledgements

This work is supported by: Birla Institute of Science and Technology (BITS) Pilani, Hyderabad campus through a research grant to G.M. (BITS/GAU/ACRG/2019/H0576) and an Institute Fellowship to G.P.S. Science and Engineering Research Board (SERB), Government of India through Ramanujan Fellowship (SB-S2-RJN-062-2017) and CRG (CRG/2020/002855) research grants to GM, and internship to N.V.P. Department of Biotechnology through research grant (BT/PR38410/GET/119/310/2020) to GM and through it, Junior Research Fellowship to R.T. We profusely thank members of the Pikaard lab for their critique of the manuscript.

## Competing interests

The authors declare no competing interests.

## Author contributions

N.V.P, R.T., and G.M. planned and designed the research; N.V.P., R.T., G.P.S., and G.M. performed the research; N.V.P, R.T., G.P.S., and G.M. analysed the data; N.V.P, R.T., and G.M. wrote the paper.

## Data availability

For this article we analyzed the publicly available/published data, and the details are as follows: The whole genome bisulfite sequencing data of *Arabidopsis thaliana* (WT and mutants) were published by Stroud et al. (2013) (GEO accession# GSM980986). The fully assembled NOR sequences of *Arabidopsis thaliana* (strain Col-0) were published by Fultz et al. (2023) and the sequences are available at GenBank as accessions OR453402 (https://ncbi.nlm.nih.gov/nuccore/OR453402) for *NOR2* and OR453401 (https://ncbi.nlm.nih.gov/nuccore/OR453401) for *NOR4*. Assembled whole genome sequences of tomato (*Solanum lycopersicum*, cultivar Micro-Tom) have been published on a preprint server (https://doi.org/10.1101/2023.10.26.564283) and the sequences are available at DDBJ (accession numbers AP028935–1 AP028946) and KaTomicsDB (https://www.kazusa.or.jp/tomato).

## Supplementary figure legends

**Figure S1:** Disruption of NOR2 gene silencing to varying degrees in DNA methylation mutants. The gel images show RT-PCR expression data for the rRNA gene subtypes and actin.

**Figure S2:** The 45S rRNA gene reference sequence used to generate methylation heat maps.

**Figure S3:** Occurrence of cytosines in the promoters and gene bodies of 45S rRNA genes. Diagram depicting the relative distribution of cytosines in CG, CHG, and CHH contexts in the promoters and gene bodies of 45S rRNA genes in (A) *Arabidopsis thaliana*, (B) tomato, and in (C) human.

**Figure S4:** Primers/oligos used in this study.

## Notes

### Competing Interest Statement

The authors have declared no competing interest.

### Summary of Updates

Some data have been removed and some data have been added based on which a new model has been proposed. Accordingly author composition has been changed.

