## Supplementary Figures for "Combination of CG methylation and Chromomethylase 2-mediated RdDM-independent CHH methylation is required for chromosome-specific rRNA gene silencing"

### rRNA subtype expression in DNA methylation mutants

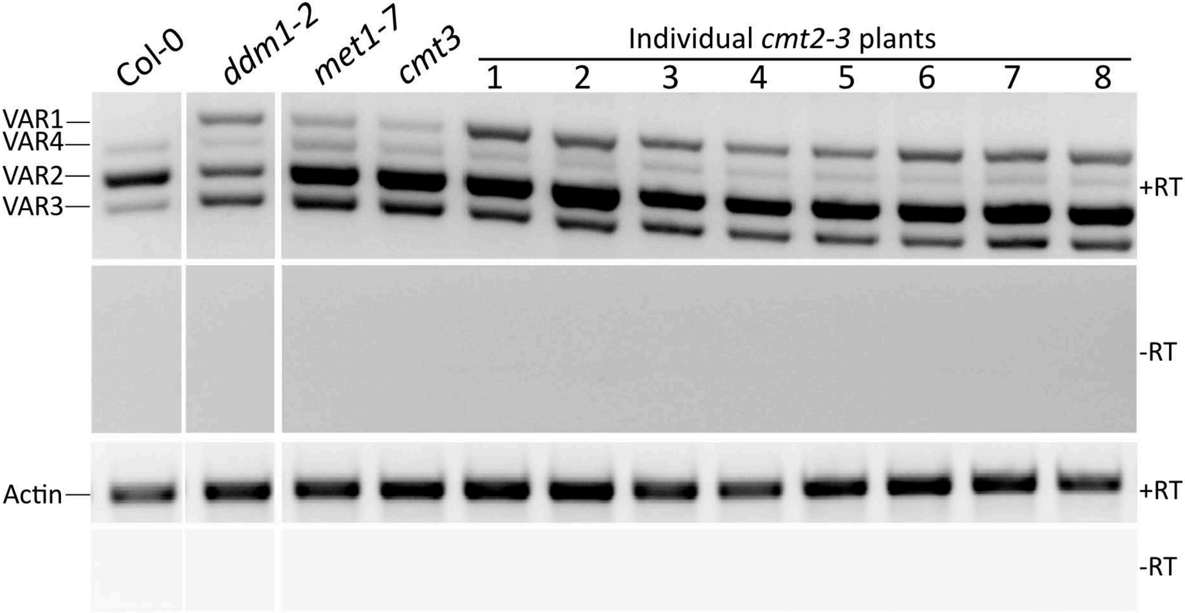


**Figure S1. Disruption of NOR2 gene silencing to varying degrees in DNA methylation mutants**

The gel images show RT-PCR expression data for the rRNA gene subtypes and actin. The table below shows the ratio of pixel intensities of VAR1:actin from the RT-PCR gel images.

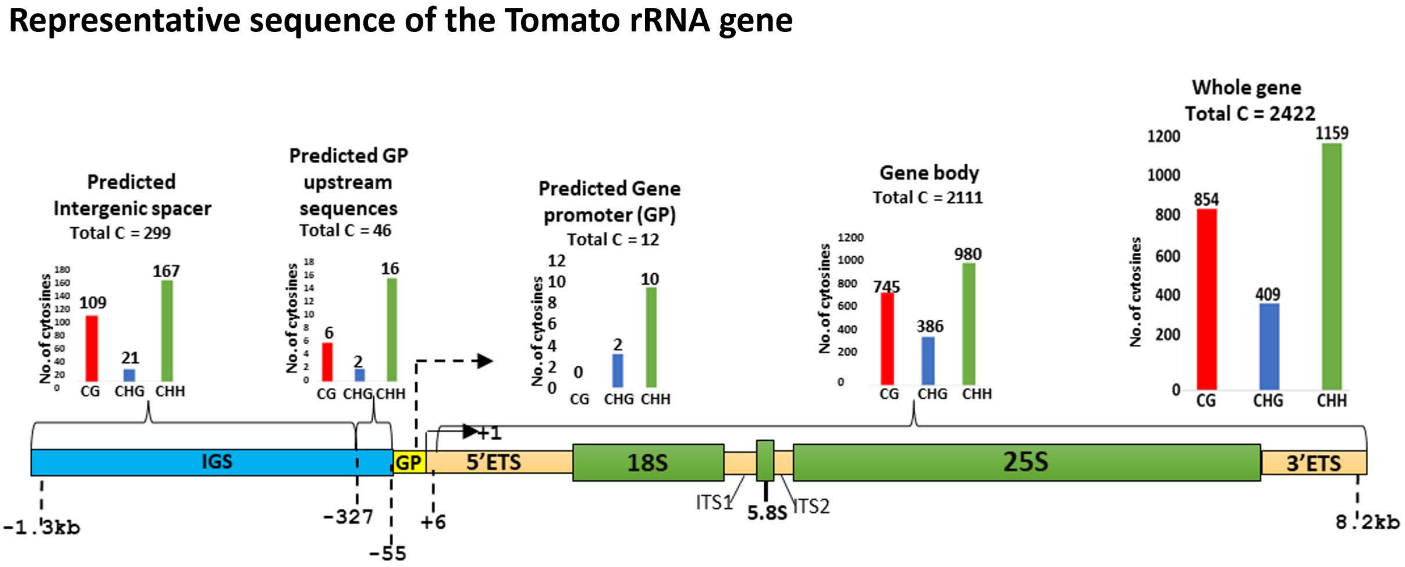


**Figure S2. Within the 45S rRNA genes, cytosine occurrence is the highest in CHH context, followed by CG, and the lowest in CHG context.** A diagram provides cytosine distribution within different regions of 45S rRNA gene sequences. Tomato NOR sequences were extracted from Shirasawa and Ariizumi, 2024. Among them, one of the gene sequences was chosen for estimating and depicting cytosine distribution in a typical 45S rRNA gene.

### Occurrence of cytosines among CG, CHG, and CHH sequence contexts in the human rDNA promoter region

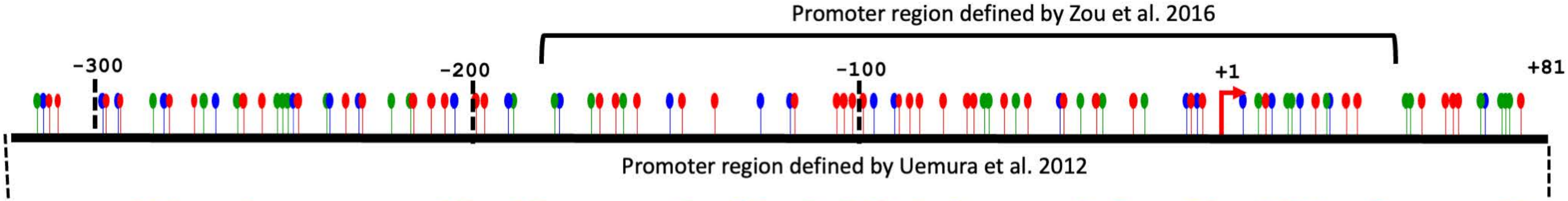


GTGTGTCCCGGTCGTAGGAGGGGCCGGCCGAAAATGCTTCCGGCTCCCGCTCTGGAGACACGGGCCGGCCCCTGCGTGTGGCC
AGGGCGGCCGGGAGGGCTCCCCGGCCCGGCGCTGTCCCCGCGTGTGTCCTGGGGTTGACCAGAGGGCCCCGGGCGCTCCGTGT
GTGGCTGCGATGGTGGCGTTTTTGGGGACAGGTGTCCGTGTCGCGCGTCGCCTGGGCCGGCGGCGTGGTCGGTGACGCGACCT
CCCGGCCCCGGGGGAGGTATATCTTTCGCTCCGAGTCGGCATTTTGGGCCGCCGGGTTATTGCTGACACGCTGTCCTCTGGCG
ACCTGTCGCTGGAGAGGTTGGGCCTCCGGATGCGCGCGGGGCTCTGGCCTCACGGTGAC

Guanosine nucleotide indicated in red font and underlined denotes the transcription start site (+1)

| Cytosine sequence context | Number of cytosines (Zhou et al., 2016) (+47 to -182) | | Number of cytosines (Uemura et al., 2012) (+81 to -310) | | Number of cytosines in the gene body (GenBank Accession U13369.1) | |
|---|---|---|---|---|---|---|
| | Number | Percentage | Number | Percentage | Number | Percentage |
| CG | 26 | ~40 | 47 | ~38 | 1870 | 39.4 |
| CHG | 19 | ~29 | 36 | ~29 | 1014 | 21.4 |
| CHH | 20 | ~31 | 40 | ~33 | 1862 | 39.2 |

**Figure S3: Occurrence of cytosines in the promoters of 45S rRNA genes.**
Diagram depicting relative distribution of cytosines in CG, CHG, and CHH contexts in the promoters of human 45S rRNA genes.

| Primer name | Sequence (5'-3') |
|---|---|
| 5'allvar | GACAGACTTGTCCAAAACGCCCACC |
| 3'allvar | CTGGTCGAGGAATCCTGGACGATT |
| Actin-F | AAGTCATAACCATCGGAGCTG |
| Actin-R | ACCAGATAAGACAAGACACAC |
| p34 | GCCGATATCCGATACCATCCCTCGATCGCTA |
| VAR1&2-R | CGATTTCCGCAACAATCACC |
| 5'-Ez-Tn5 | CTGTCTCTTATACACATCTGGTTACCAGGGCCCAGCGGCCGCAGATGTGTATAAGAGACAG |
| 3'-Ez-Tn5 | CTGTCTCTTATACACATCTGCGGCCGCTGGGCCCTGGTAACCAGATGTGTATAAGAGACAG |

**Figure S4: Primers/oligos used in this study.**
